# High fidelity epigenetic inheritance: Information theoretic model predicts *k*-threshold filling of histone modifications post replication

**DOI:** 10.1101/2021.05.25.445560

**Authors:** Nithya Ramakrishnan, Sibi Raj B Pillai, Ranjith Padinhateeri

## Abstract

Beyond the genetic code, there is another layer of information encoded as chemical modifications on histone proteins positioned along the DNA. Maintaining these modifications is crucial for survival and identity of cells. How the information encoded in the histone marks gets inherited, given that only half the parental nucleosomes are transferred to each daughter chromatin, is a puzzle. Mapping DNA replication and reconstruction of modifications to equivalent problems in communication of information, we ask how well enzymes can recover the parental modifications, if they were ideal computing machines. Studying a parameter regime where realistic enzymes can function, our analysis predicts that, pragmatically, enzymes may implement a threshold − *k* filling algorithm which fills unmodified regions of length at most *k*. This algorithm, motivated from communication theory, is derived from the maximum à posteriori probability (MAP) decoding which identifies the most probable modification sequence based on available observations. Simulations using our method produce modification patterns similar to what has been observed in recent experiments. We also show that our results can be naturally extended to explain inheritance of spatially distinct antagonistic modifications.

## INTRODUCTION

Why do our bone cells behave very differently from our muscle cells or cells of other types even though they all have the same genetic code? To explain the emergence of such diverse cell types, one would need to note that, beyond the genetic code, there are multiple layers of information encoded by the wrapping and folding of the DNA into chromatin with the help of many proteins [1, 2]. Most of the DNA is wrapped around octamers of histone proteins, making the chromatin, essentially, like a string of beads made of nucleosomes (DNA+histones) [3–6]. Each nucleosome carries chemical modifications, like acetylations and methylations [7], forming a pattern of histone marks along the chromatin polymer contour [2, 8–10] (see FIG. 1(A) top panel). This pattern encodes a crucial layer of information regulating accessibility, transcription, replication and other cellular processes. Even though the entire histone modification code is not deciphered yet, we understand it in parts. For example, H3K27me3 represses reading of the local DNA region where the modification is present, H3K9ac encodes for local gene activation and so on [7, 11, 12]. The activation and repression of genes collectively decide the gene expression pattern and hence determine the function and fate of a cell [10, 13, 14, 14, 15].

**FIG. 1:**
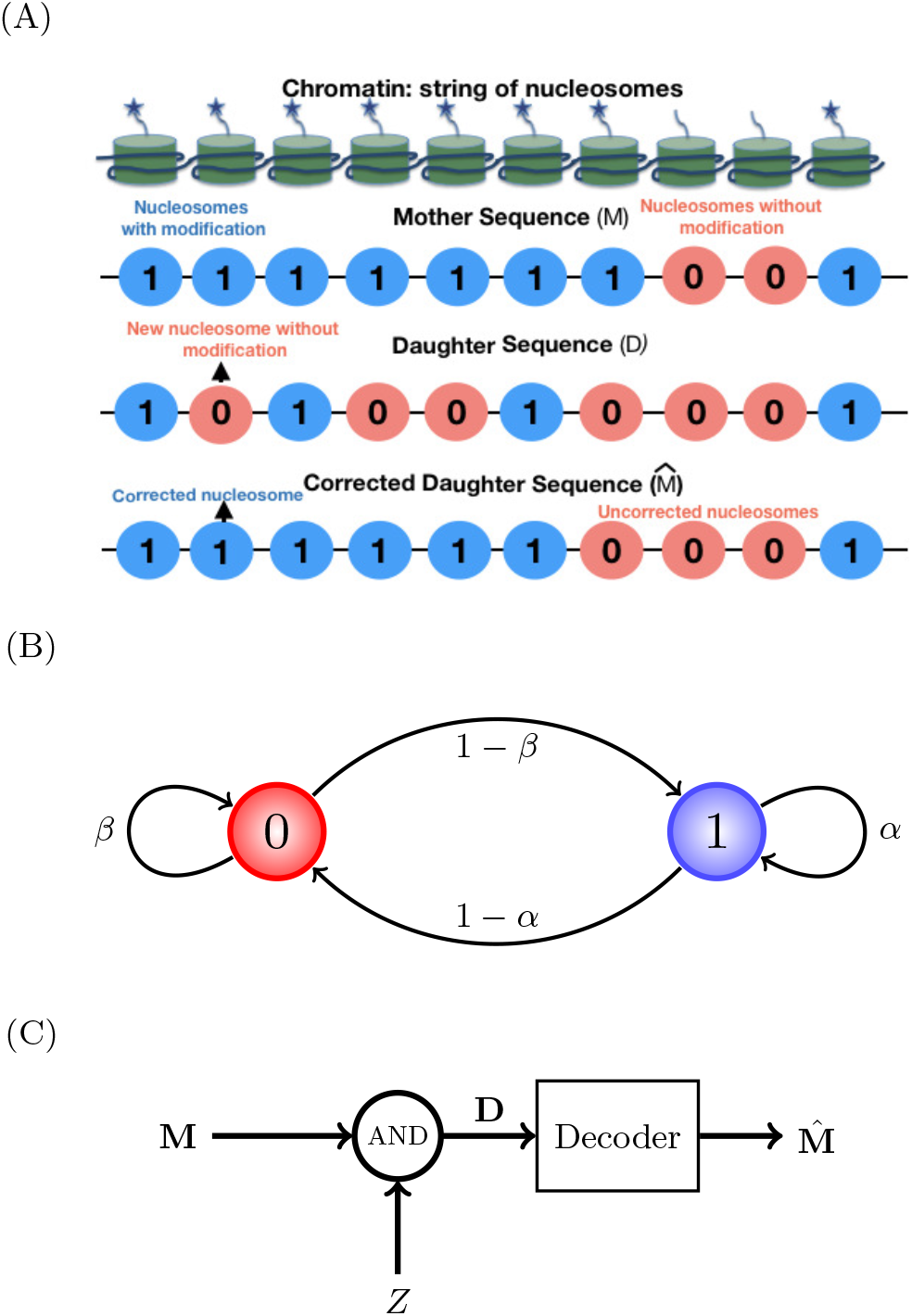
Schematic description of the problem (A) Row 1 from top: chromatin as a string of nucleosomes with and without the histone modification (star) of our interest. This can be mapped to a string of binary numbers indicating the presence (1) or absence (0) of the modification (row 2), giving us **M**. Row 3: one typical realization of a daughter chromatin (**D**) produced from **M** above, via a process mimicking DNA replication, where only a fraction of the modifications (1s) will end up in the daughter chromatin, stochastically; the rest do not have the modification of interest (0) [24]. Soon after replication, certain enzymes will insert modifications correcting **D** to a mother-like sequence 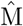 (row 4). Since these are stochastic processes, we expect some errors. (B) The mother sequence (**M**) is modeled as a first order Markov chain having sequence of 0s and 1s. *α* and *β* are probabilities of finding a 1 followed by a 1, and a 0 followed by a 0, respectively. 1 − *α* and 1 − *β* are probabilities of finding a 1 followed by a 0 (note arrowheads), and a 0 followed by a 1, respectively. (C) The daughter sequence (**D**) is obtained by a mother sequence **M** getting logically ANDed with an independent and identically distributed (IID) binary sequence **Z** (noise). A mother-like sequence 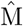 is reconstructed by passing **D** through a decoder. The plausible ways by which enzymes could act as decoders is the subject of this study.

While preparing to divide, cells copy their genetic code via the DNA replication process. For DNA to be copied, the chromatin has to be unfolded and histone proteins need to be disassembled [16, 17]. This would disrupt the pattern of histone modifications. Recent studies have shown that the (H3 − H4)_2_ tetramer from the parent remains intact [18] and randomly gets deposited onto either of the newly synthesized DNA strands [19–21]. That is, daughter strands will have only some (≈ 50%) of its nucleosomes from the parent; the rest need to be assembled from the pool of new histone proteins made afresh [22, 23]. Since the new nucleosomes will not carry the histone marks present on the parental chromatin, half the parental marks are missing in each daughter chromatin [24]. Since the histone modification patterns can decide the state (repressed/active) of all genes and the identity of the cell itself, it is crucial that the newly made chromatin re-establishes the pattern immediately after replication [25]. Recent experiments show that many of the histone modification patterns – patterns in the repressed gene regions, in particular – are “inherited” from the parental chromatin [26–28]. How the new chromatin recovers the missing information and re-establishes a mother-like pattern is a puzzle.

It is known that there are specific enzymes to read and write histone marks [29, 30]. While molecular details of some of these enzymes are known [31–33], precisely what strategies they use to re-establish the histone modification pattern after replication are not fully understood. Previous theoretical studies have investigated average properties of modifications and whether long-range interactions are necessary [34] or short-range interactions would suffice [35]. Additionally, the bivalent and bistable states of the chromatin modifications have been explored theoretically and supported by experimental observations [36–38]. There also exist models that discuss the establishment of modifications in clusters or as a steady state in the inherited chromatids [39–44]; however, the experimentally observed high-fidelity re-establishment of the modification patterns [26] is not explained by the existing models.

The problem of loss of information in the modification pattern during replication and its retrieval within the daughter chromatin is very similar to data loss and error correction in telecommunication. In such systems, a transmitted signal gets exposed to noise and consequently becomes, error-prone at the receiving end. The decoder at the receiver detects and corrects these errors using techniques from information and coding theory [45, 46]. This viewpoint immediately poses the following questions: Can we use known decoding algorithms from communication theory to analyze chromatin modification loss and retrieval? How well can the best known algorithms correct the missing modifications and re-establish the modification patterns? What is the best possible correction strategy enzymes could use if they were ideal computing machines? Are these algorithms compatible with the biological processes that realistic cellular enzymes can conceivably do? In this paper, we address these questions using ideas from Information theory. We consider one of the daughter chromatins to be a noise-corrupted signal created at the replication fork, while the enzymes and other molecular agents help to correct this error using mathematical techniques. In this model, the inheritance of the mother’s pattern is approached using Bayesian decoding techniques. Predictions from our model are verified using publicly available experimental data, indicating the relevance of our work in studying real biological datasets [26].

## MODEL AND METHODS

Consider a region on a mother chromatin having *N* nucleosomes. We are interested in studying the inheritance of one histone modification at a time. Since many of the repressive marks are known to be inherited accurately [26] after replication, we will consider one such repressive mark (e.g., H3K27me3) and its pattern along a chromatin. This pattern can be represented by a vector **M** = {*m*_1_, *m*_2_, …, *m*_*N*_}, where *m*_*i*_ can have values 1 or 0 indicating the presence or absence of the modification on the *i*^*th*^ nucleosome (see FIG. 1(A)).

For brevity, we use the following notations:

- 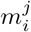 represents the sequence of modification values (*m*_*i*_, *m*_*i*+1_, …, *m*_*j*_) between location *i* and *j* with *j* > *i*; Thus, 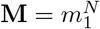 represents the entire mother chromatin modification sequence.
- a region with *k* consecutive modifications present (ones) or absent (zeros) will be denoted as 1_*k*_ or (0_*k*_), respectively. Extending this, the sequence (1, 0, …, 0, 1) representing an island of *k* consecutive zeros between two ones will be denoted as (1, 0_*k*_, 1).

Since modification on a nucleosome is very likely related to its immediate neighbors, we model the pattern **M** along the mother chromatin as a binary-valued random walk, having neighbourhood interactions corresponding to a first order homogeneous Markov chain. More specifically, given the modifications *m*_*i*−1_ and *m*_*i*+1_, the modification *m*_*i*_ is assumed to be independent of all other modification values. Equivalently, the conditional probability law is

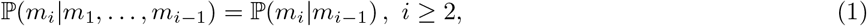

where *m*_1_ is the modification on the first nucleosome of the region of our interest. The state-space evolution of the Markov chain **M** is as follows: given *m*_*i*_ = 1, let *α* and 1 − *α* be the probabilities for obtaining *m*_*i*+1_ = 1 and *m*_*i*+1_ = 0 respectively. Similarly, if *m*_*i*_ = 0, let *β* and 1 − *β* be the probabilities for having *m*_*i*+1_ = 0 and *m*_*i*+1_ = 1 respectively. The sequence **M** can be seen as a random walk on the state space shown in FIG. 1(B). The parameters *α* and *β* are functions of the mean contiguous length of the modified and unmodified regions respectively and can be computed from the experimental data. See Supplementary Information (SI). For example, when *α* and *β* values are close to 1, the pattern would often contain long runs of either 1s (presence of modification) or 0s (absence of modification).

From the mother chromatin **M**, the generation of a daughter chromatin having histone modification sequence 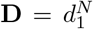 is modeled as follows. During replication, with probability 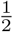, each nucleosome on a daughter chromatin is either directly inherited from its parental counterpart (i.e. *d*_*i*_ = *m*_*i*_) or newly deposited (i.e. *d*_*i*_ = 0) from a pool of fresh histones assembled de novo [24]. We consider both the histones in the (*H*3 − *H*4)_2_ tetramer to be symmetrically modified in our model based on recent evidence that a large proportion of the histones are symmetrically modified [47, 48]. This process is equivalent to doing a logical AND operation of the mother sequence **M** with an independent binary vector **Z** (noise), which is generated by independent tosses of a fair coin. Thus, **D** = **M**.**Z** (see FIG. 1(C)), where **Z** has Independent and Identically Distributed (IID) entries. This biological process leads to a memoryless model with the conditional probability

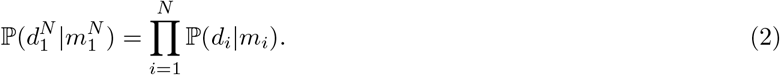

Owing to the independent and random nature of the placement of the parental nucleosomes along the newly formed daughter chromatin, the above relation is justified. In biology, the question is, given a daughter sequence 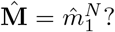 soon after replication, how can a cell reconstruct a mother-like sequence 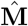 In other words, is it possible to build a decoder that would reconstruct a 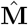 from **D**, as depicted in FIG. 1(C)? Ideally, a cell would want to choose a binary sequence 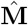 having the minimum deviation from **M**. A simple way to quantify this deviation is to compute the fraction of errors in the reconstructed sequence as:

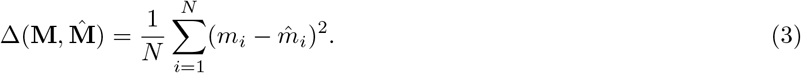

Since we are comparing bitwise, this deviation metric is effectively the bit error rate (BER) when *N* becomes large [49]. In communication, BER is the proportion of bits that are in error among the total transmitted/received bits. In the present context, BER is essentially the fraction of nucleosomes in a daughter sequence that differ in their histone mark from the parental sequence. Thus the chosen 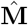 should minimize the BER with respect to the actual sequence **M**, while obeying the transition law in Eq. (1). This is similar to data communication through an erroneous channel. It is well known that Bayesian estimation schemes minimize the average detection error probability at the receiver. In particular, a decoder choosing the input sequence having the Maximum Àposteriori Probability (MAP) is optimal in minimizing the message error probability in communication [46, 49]. We call this the Sequence MAP (SMAP) decoder, which identifies the most probable sequence 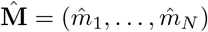 based on the observations 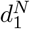 as

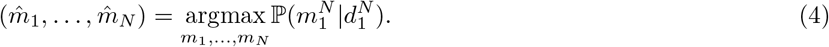

The above equation essentially says that, given a daughter sequence, the sequence that maximizes the conditional probability should be declared as the reconstructed mother-like sequence. SMAP decoding is known to have very good BER performance and good analytical tractability in many contexts [49]. While the optimal BER performance can be achieved by Bitwise MAP (BMAP) decoding for each modification value separately, the latter scheme is not only computationally more demanding, but also analytically less tractable. It appears that SMAP decoding is a potential candidate for biological cells to reconstruct the epigenetic modifications from the partial data. Notice that SMAP decoding depends on the parameters *α* and *β* of the Markov chain. Clearly, this algorithm is only targeting a primary reconstruction immediately following DNA replication; secondary mechanisms like synergistic activities of correlated modifications or DNA methylation maintenance post replication over the course of the cell cycle may further alter the patterns in the long run [19, 24, 50–52].

## RESULTS

In this section, we will be using the ideas developed in the Model and Methods to answer how one can reconstruct a mother-like modification sequence 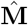, given the daughter chromatin sequence **D**. We will discuss how well algorithms like the Sequence MAP (SMAP) decoding will compute 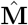, and whether realistic enzymes can implement this in practice.

## Ideal Enzymes Implementing SMAP Decoding

While we do not know exactly how biological enzymes work to retrieve histone modification patterns soon after replication, how well can the communication theory-inspired algorithms reconstruct a mother-like sequence? To test this, one can imagine some *ideal enzymes*—computing machines—constructed to implement the SMAP algorithm in Eq. (4). That is, these enzymes will maximize the conditional probability ℙ (**M** | **D**) over possible sequences of **M**, based on the given daughter sequence observations 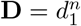. Applying Bayes’ rule [53], along with Eq. (1) and Eq. (2) (see Supplementary Information (SI) Sec. I), one gets

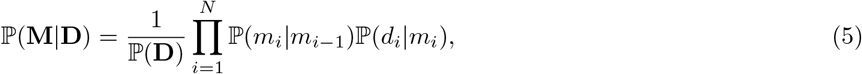

where we took *m*_0_ = ∅ (empty set) for notational convenience. Since ℙ (**D**) does not play a role in the maximization over **M**, it can be ignored for our purposes. The equation says that the reconstruction process of the mother-like sequence needs to account two facts: the modification status of the immediate neighbours (ℙ (*m*_*i*_ | *m*_*i*−1_)) and the information embedded in the replication process (ℙ (*d*_*i*_ | *m*_*i*_)). Knowing Eq. (5), ideal computing machines can now implement the SMAP algorithm using the idea of trellis decoding [45], which is closely related to the well known Viterbi Algorithm in coding theory [49]. While the memory and computational power requirement for trellis decoding is high in general, we find that decoding procedure for our model can be broken down to smaller sub-sequences, each corresponding to a different run of zeros in **D**. In particular, SMAP decoding can be applied separately on sub-sequences of the form (1, 0_*k*_, 1), which has *k* consecutive zeros in between two ones. Mathematically, we can show that SMAP decoding will choose a sequence having 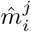 according to 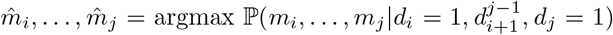; here *i* and *j* be two positions (with *j* > *i*) where the daughter sequence has ones (see SI Sec. III). Our analysis suggests that to decide on a bit at position *l* where the daughter has inherited a zero, we need to only consider the smallest daughter segment containing the position *l*, and flanked by ones at both ends. Only daughter segments with at least one intermediate zero are to be considered; otherwise there is nothing to decode. Without loss of generality, we will take **D** = (1, 0_*k*_, 1) for the rest of the exposition, corresponding to a run of *k* zeros, and perform trellis decoding on this sequence.

### Trellis Decoding

Trellis decoding [45] is a technique that can identify the most probable mother-like histone modification sequence, according to the theory described above. In this study, a sequence of states of a Markov chain (here, histone modification sequence) is called a path or a trajectory. Given the states at the start and end of a possible path, a trellis diagram [45] can be used to find the joint probability of each path with the given observation **D**. Since **D** has the form (1, 0_*k*_, 1), the start and end states of the trellis are ones. FIG. 2 demonstrates the trellis diagram for a run of 5 zeros (*k* = 5). Starting from the initial state 1 (left bottom in FIG. 2), the trellis diagram assigns a conditional probability (branch metric) to each subsequent transition (arrow), based on the transition probability law and observed daughter state. For transitions from state at *i* − 1 to *i*, the branch metric is ℙ (*m*_*i*_, *d*_*i*_|*m*_*i*−1_), which can be evaluated as ℙ (*m*_*i*_ | *m*_*i*−1_)ℙ (*d*_*i*_ *m*_*i*_), where *d*_*i*_ = 0 for 2 ≤ *i* ≤ *N* − 1. While the sequence **D** is easily seen to be generated by a hidden Markov Model (HMM), the branch probability metrics are explicitly given inside the box of FIG. 2 (also see SI Sec. II). Notice that we have to find ℙ (*m*_*i*_, *d*_*i*_ = 0|*m*_*i*−1_) for (*m*_*i*−1_, *m*_*i*_) *∈ {*(0, 0), (0, 1), (1, 0), (1, 1)*}*, and ℙ (*m*_*i*_ = 1, *d*_*i*_ = 1|*m*_*i*−1_) for *m*_*i*−1_ *∈ {*0, 1*}* for the last stage (see FIG. 2).

**FIG. 2:**
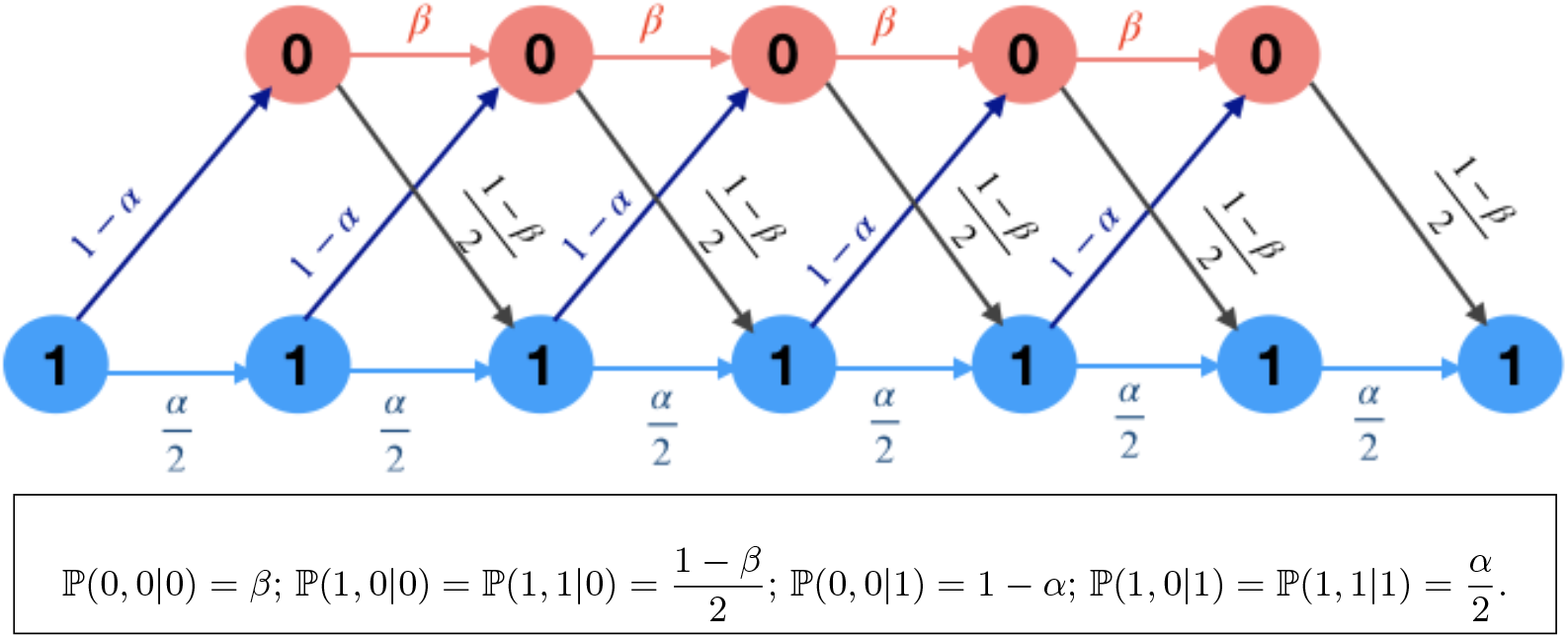
The trellis diagram used for illustrating how the MAP algorithm chooses the path of maximum a posteriori probability given a daughter sequence (1, 0_5_, 1). From each Markov state (0 or 1), there are two possible arrows to transition to the next state. The values ℙ (*m*_*i*_, *d*_*i*_ | *m*_*i*−1_) given in the box represent probabilities associated with each arrow. Starting from state 1 at the left bottom, for each possible path moving along the arrows, one can compute the path probabilities (path metrics) using Eq. (5). We finally choose the path with the largest path metric.

Notice that each possible mother sequence can be identified as a path in the trellis, with the labels identifying the branch metrics. Using Eq. (5), the product of corresponding branch metrics will yield the path metric of each possible sequence, and then the path maximizing the SMAP metric can be chosen.

On a computer, we generated several mother sequences for different values of *α* and *β* parameters in the Markov model. We then generated several daughter sequences, for each of the mother sequences, by flipping 1s to 0s randomly with probability 0.5; that is, **D** = **M · Z**, with **Z** an IID binary sequence generated by an unbiased coin. Each of the daughter sequences was corrected with the trellis-based SMAP decoding to generate the corresponding estimate 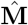; the error 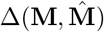 was computed (Eq. (3)). The average of 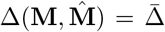 (averaged over many realizations) for fixed pairs of *α* and *β*, is presented as a heatmap in FIG. 3. The mean deviation 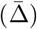 between mother and corrected mother-like daughter is low for very high values of *α* and *β* — a region dominated by long islands of ones (modified nucleosomes) separated by long islands of zeros (unmodified nucleosomes). There are other regions too where 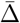 is relatively small (also see SI Sec. III, FIG. S1). Overall, this result shows how well an ideal computing machine that employs state of the art Information theory inspired algorithms can recover the original mother sequence. The remaining question is, can a real enzyme do as good as this computing algorithm?

**FIG. 3:**
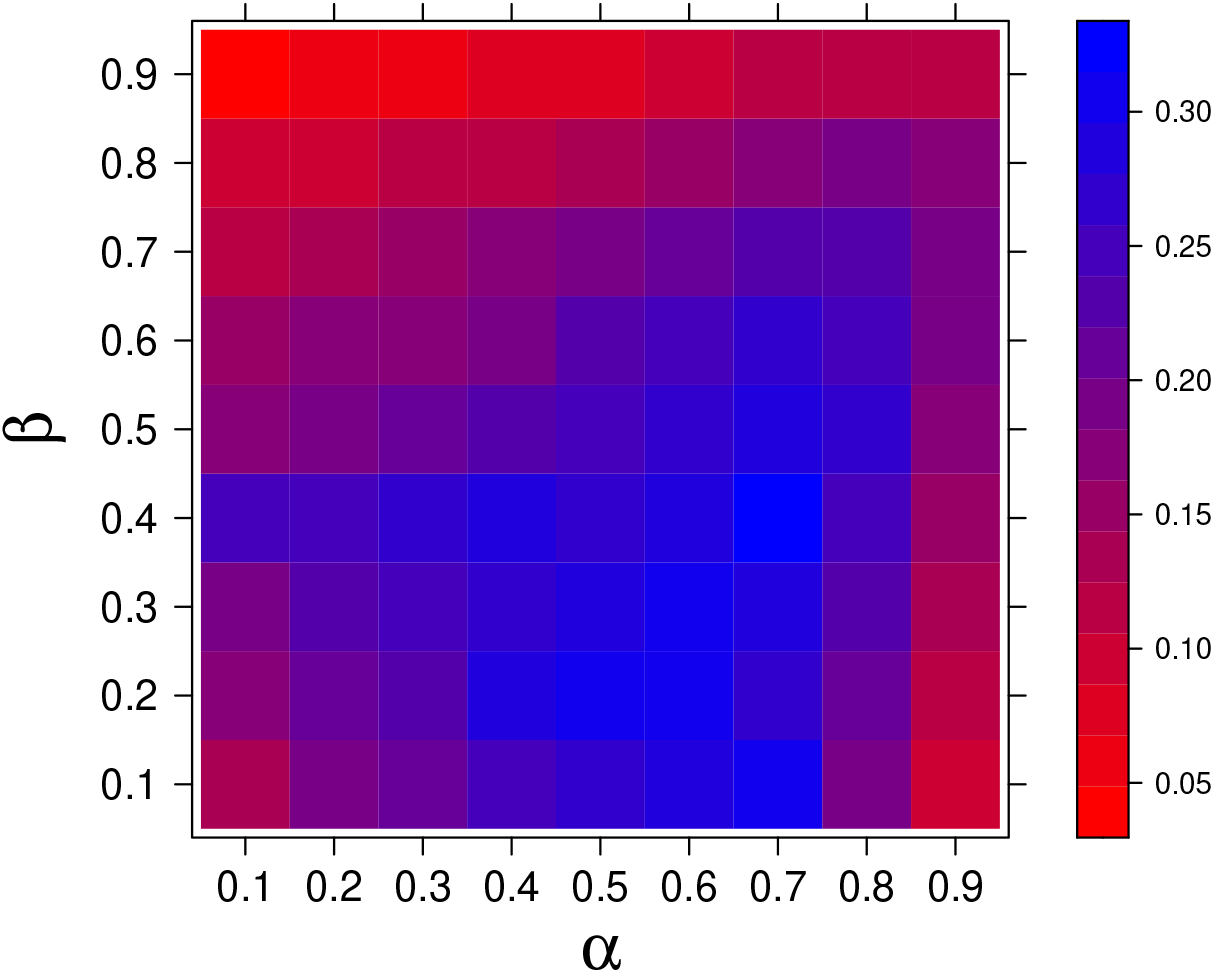
The average deviation between the original mother and the mother-like corrected daughter sequences (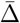 = ensemble averaged 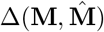) is plotted for different *α* and *β* values as a heatmap (see color bar on the side). The error is averaged over the error 300 mother sequences and 200 daughter sequences corresponding to each mother sequence — that is, 60000 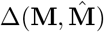 values.

## Threshold-*k* model: enzymes filling unmodified islands of size at most *k* maintain chromatin fidelity

Whether biological enzymes are equipped to do complex SMAP computations like trellis decoding by themselves is debatable. Nevertheless, we argue that in certain biologically relevant parameter regimes, the decoding rule can be simple enough for enzymes to potentially execute. Among the known histone modification patterns, it is common to have regions densely filled by a certain modification (e.g., H3K27me3), and other regions where the modification is totally absent. This corresponds to higher values of *α* and *β* in our Markov model (see FIG. 1(B); also see SI Sec. IV). Below we show that in this regime, the SMAP algorithm simplifies to tasks that the enzymes may easily carry out.

Consider an island of *k* unmodified nucleosomes in the daughter chromatin, giving the pattern (1, 0_*k*_, 1). From the trellis diagram (FIG. 2), it can be seen that, if 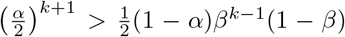, the probability of having the all-one path (1, 1_*k*_, 1) at the mother is greater than that of (1, 0_*k*_, 1). In other words, when the function 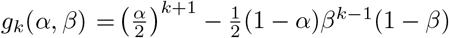 is positive, an all-ones path is preferred over a run of *k* zeros by SMAP decoding. We can characterize the values of *α* and *β* for which the above condition holds true. The expression *g*_*k*_(*α, β*) = 0 is easy to solve if we take *k* to be a real value, this yields the root *k*^*^ as:

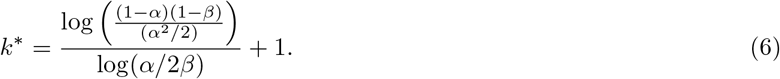

Notice that the solution for *k*^*^ is unique when 0 < *α* < 1 and 0 < *β* < 1. Since 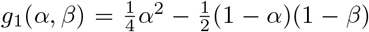, the condition *g*_1_(*α, β*) > 0 implies that the sequence (1, 1, 1) is preferred over (1, 0, 1). In addition, this condition will also imply that the numerator of Eq. (6) is positive. The uniqueness of *k*^*^ will now have the following implications:

On a unit square region of parameters (*α, β*) *∈* (0, 1) × (0, 1),

- When *α* < 2*β* and *g*_1_(*α, β*) > 0, one gets *k*^*^ *≥* 1; we find that *g*_*k*_(*α, β*) > 0 for all positive integers *k ≤ k*^*^ in Eq. (6). This suggests that the SMAP algorithm will replace (1, 0_*k*_, 1) with (1, 1_*k*_, 1), if and only if *k ≤ k*^*^.
- When *α* > 2*β* and *g*_1_(*α, β*) > 0, we find that *g*_*k*_(*α, β*) > 0 for any positive integer *k*; hence the SMAP algorithm will replace (1, 0_*k*_, 1) with (1, 1_*k*_, 1), for any value of *k*.

Notice that when every path of at most *k*^*^ zeros between two ones has less path metric than the corresponding all ones path, clearly any possible path other than all ones cannot have the maximum SMAP metric, while decoding sequences of length less than *k*^*^.

The above analysis based on trellis decoding suggests two simple ways for enzymes to work. Enzymes of Type-*I* would simply modify all unmodified nucleosomes (0s) between two modified nucleosomes (1s). Such enzymes may be preferred when the modification pattern can be modeled by parameters *α* and *β* that corresponds to region *a* in FIG. 4(A); notice that this has large *α* and small *β*. An enzyme of Type-*II* would fill all unmodified nucleosomes (0s), if and only if the size of the unmodified region is ≤ *k*^*^. That is, replace (1, 0_*k*_, 1) by (1, 1_*k*_, 1), if the island size is *k* ≤ *k*^*^. Thus long islands of 0*s* are left unfilled. We call this a **threshold-***k* **filling model**, which becomes active in region *b* of FIG. 4(A). Notice that when both *α* and *β* are close to 1, the modification is expected to have long domains (islands) with its presence, followed by islands with no modification. Biologically, this is a realistic regime for many modifications where enzymes can do threshold-*k* filling.

**FIG. 4:**
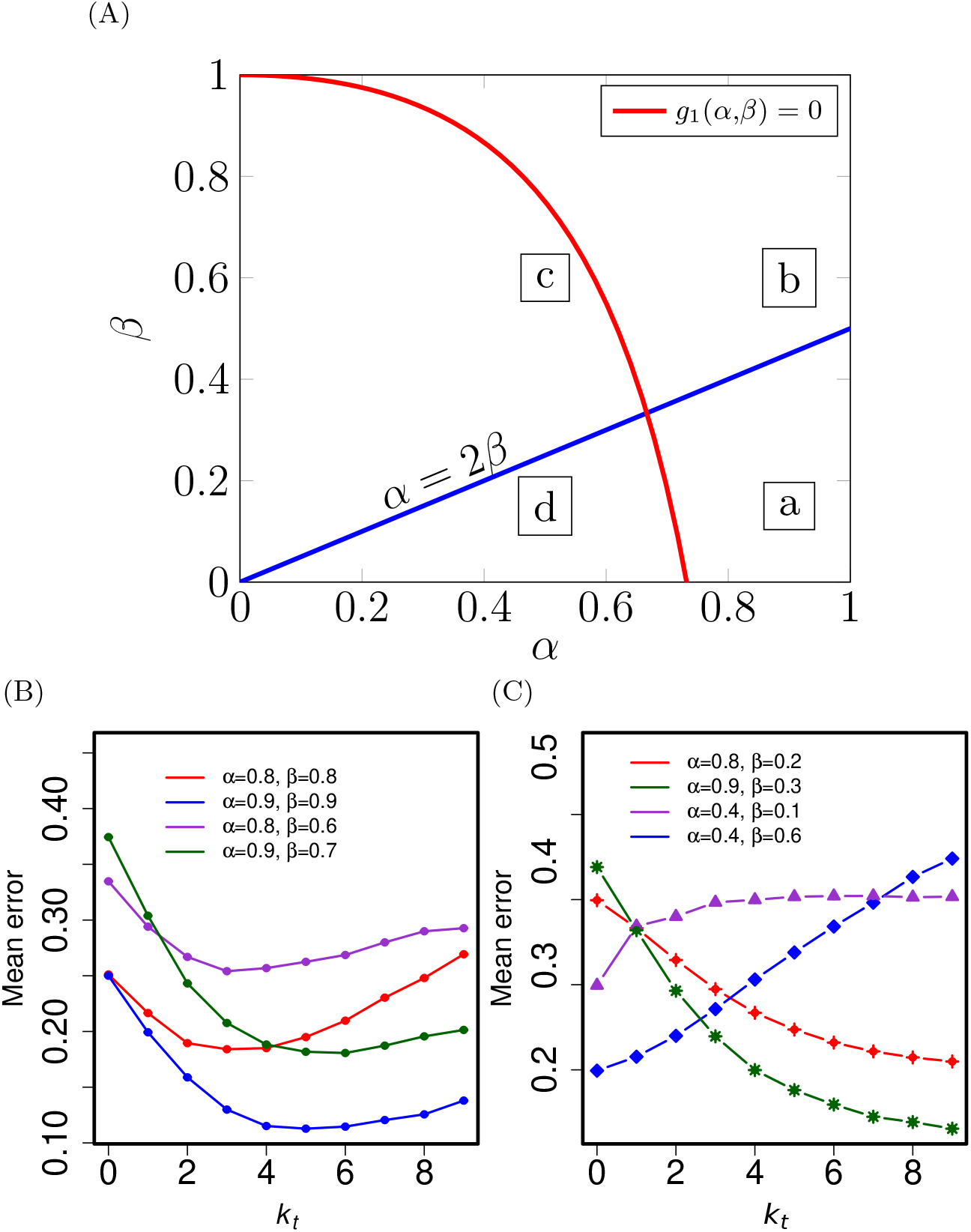
Behavior of our algorithms across the parameter spaces (A) Four (*a, b, c, d*) regions in (*α, β*) parameter space. The curves shown are *g*_1_(*α, β*) = 0, and *α* 2*β* = 0. In region *a* (*g*_1_ > 0, *α* > 2*β*), the SMAP will replace every 0 with 1. In the region *b* (*g*_1_ > 0, *α* < 2*β*), enzymes can implement threshold-*k* filling (see text). This parameter regime is realistic, biologically. (B) The mean error 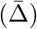 when we fill all islands of 0s having size at most *k*_*t*_. All these curves have (*α, β*) values in region *b*, and 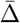 is non-monotonic having a finite optimum *k*_*t*_ = *k*^*^. (C) 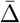 for parameter values in region *a* (red and green curves) are monotonically decreasing suggesting that the optimal *k*_*t*_ is unbounded; hence the least error would be when all 0s are replaced with 1s. In regimes *c, d* (blue and violet curves) the mean error is minimal when nothing is filled suggesting that threshold-*k* filling is not suitable here. The standard errors here are smaller than the size of the points.

We tested the threshold-*k* filling model on a computer by generating several mother and daughter sequences, for various values of *α* and *β*; each daughter sequence was corrected using the threshold-*k* filling algorithm — that is, we filled all islands of 0s, having size *k* ≤ *k*_*t*_, by 1s; here *k*_*t*_ is taken as a variable. Corresponding mean error 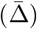, averaged over many realizations, was computed for a given (*α, β*). In FIG. 4(B), all the curves correspond to (*α, β*) in regime *b* of FIG. 4(A) (*g*_1_ > 0 and *α* < 2*β*). In this interesting regime, the mean error has a non-monotonic behaviour with a minimum at a particular value of *k*_*t*_ = *k*^*^, where *k*^*^ is predicted by Eq. (6). Note that *k*^*^ values are around 3 to : enzyme of Type-*II* can fill small unmodified islands having 3 to 6 zeros, and leave much longer unmodified islands unfilled.

In FIG. 4(C, the two monotonically decreasing curves belong to the parameter regime *a*, (i.e.) (*g*_1_ > 0, *α* > 2*β*). The mean error is decreasing as we increase *k*_*t*_, suggesting that there is no finite *k*^*^. If the modification patterns were in this regime, the corresponding enzymes should attempt to fill every unmodified region, however small or big that may be. The other two curves in FIG. 4(C) correspond to regimes *c* and *d* in FIG. 4(A). For both the curves, the mean error is increasing, suggesting that the threshold-*k* filling algorithm is not suitable in these parameter regimes. It is also unlikely that these parameter regimes would be biologically relevant.

## *k*-threshold filling model can obtain inheritance patterns similar to what is observed experimentally

To examine the biological relevance of the findings leading to a *k*-threshold filling model, we took publicly available experimental histone modification data, and compared with our simulation results. The inheritance of modification H3K27me3 has been systematically studied recently [26] by measuring modification occupancy before and after DNA replication. Since the experimental data is population-averaged, we used a simple randomized discretization algorithm to generate many binary sequences of the available data (see SI Sec. V). For example, we took a long region from the chromosome 1 (151,495,060 bp to 165,790,665 bp) data, discretized to obtain the mother vector (**M**), and then generated the daughter sequence **D** = **M** · **Z**. The **D** was corrected to a mother-like modification sequence 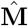 using the threshold-k algorithm for different *k* = *k*_*t*_. This was repeated several times (100 **M** sequences, and 100 **D** for each **M**), and the mean error 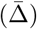 is plotted as a function of *k*_*t*_ in FIG. 5(A). The results show that when *k*_*t*_ = 5 the mean error in the corrected H3K27me3 pattern is minimum. Note that this is very similar to the curves in FIG. 4(B) for large values of *α* and *β*. We independently verified that the parameters corresponding to the original mother sequence is *α* ≈ 0.81 and *β* ≈ 0.815 (see SI Sec. VI) implying that a biologically relevant modification falls in parameter regime *b*.

**FIG. 5:**
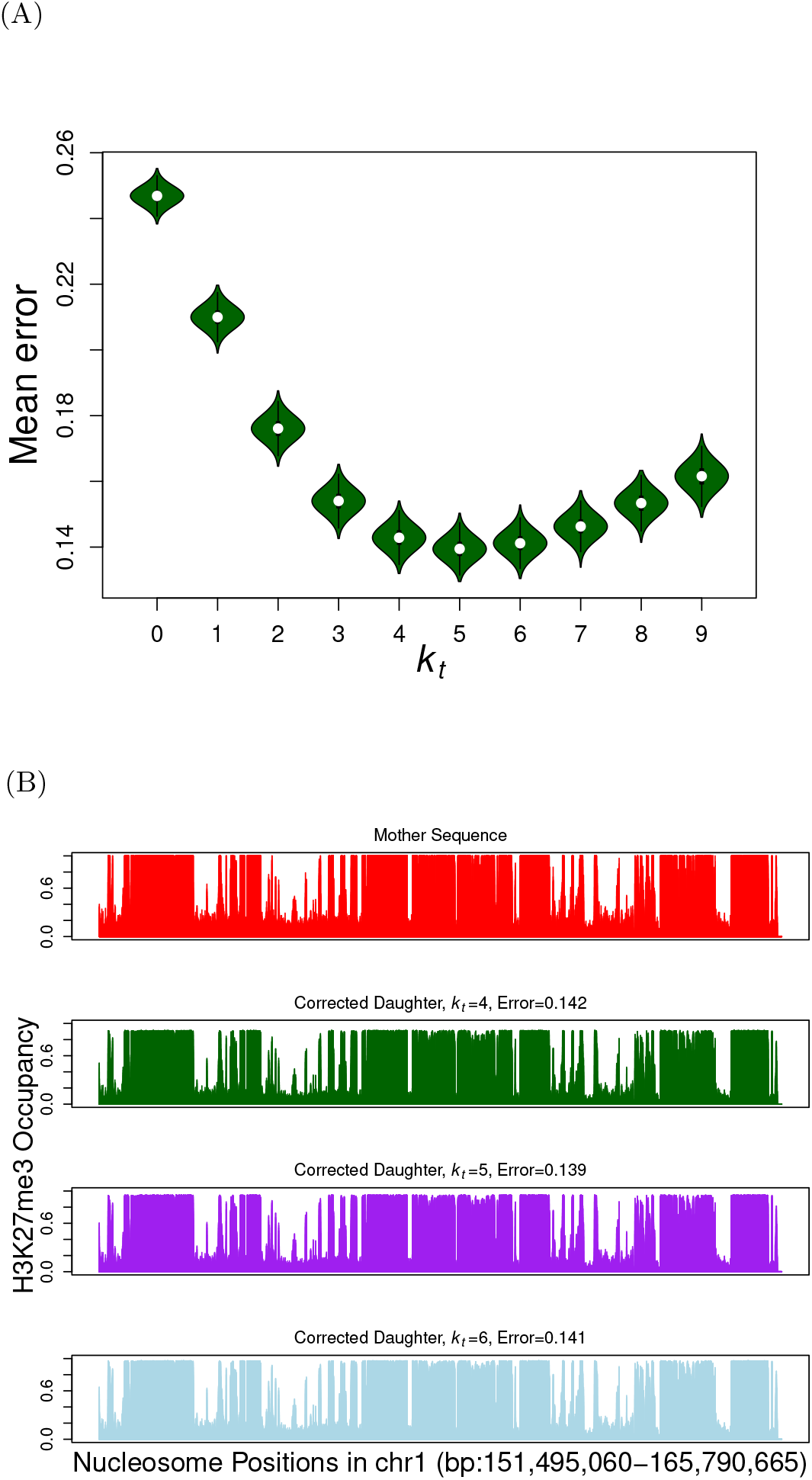
Error Correction in H3K27me3 data (a) Mean deviation 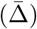 between corrected daughters and corresponding mothers, where the experimental population-averaged parental data for H3K27me3 is from [26] (see database GEO: GSE110354). Error correction was performed using the threshold-*k* filling algorithm for different *k*_*t*_ values. (b) The population averaged histone modification occupancy for H3K27me3 is plotted for mother sequence (top), and corrected daughter sequences corresponding to different values of *k*_*t*_.

In FIG. 5(B), we plot the population-averaged modification pattern for different values *k*_*t*_. Note that islands of high and low modification occupancy regions are present in both the mother as well as the corrected daughter sequences. This indicates that our threshold *k*-filling algorithm can reproduce biologically relevant data. Thus, our information theory-inspired algorithm predicts that there might be enzymes that simply fill short segments (4 or 5 nucleosomes) of unmodified regions, but leave the longer unmodified regions (> 5 nucleosomes) unfilled. This helps in maintaining the fidelity during epigenetic inheritance. The two interesting biological questions in this context, namely (i) what are the plausible biological processes/mechanisms that could facilitate such a threshold filling process and (ii) how enzymes know what is the optimal threshold for filling, are discussed further in the Discussion section.

## Spatially distinct antagonistic modifications

The threshold-*k* filling model can be naturally extended to study two (or multiple) modifications that are antagonistic, spatially distinct (the same nucleosome will not have both the modifications simultaneously), and to be acted upon by very different enzymes. In the context of the bivalent/bistable state of the chromatin discussed in [54], it has to be noted that the antagonistic model we consider is not bivalent (i.e), the antagonistic modifications are not present in the same nucleosomes simultaneously. As per our model, for an enzyme-1 responsible for modification-1, the nucleosomes having the second modification are not “visible” and would appear as a long stretch of 0s. For enzyme-2, similarly the modification-1 nucleosomes appear as a long stretch of 0s. We generated a sequence having two spatially distinct modifications (see SI Sec. VII). A simple extended version of the threshold-*k* algorithm was applied to each enzyme separately. That is, whenever there is an island of size *k* ≤ *k*_*t*_ between two nucleosomes having modification-*i*, the region was filled with modification-*i*, for *i* = 1, 2. Anything else was left unfilled. This gave us results as shown in FIG. 6. These are occupancies from a typical realization (not averaged over the population) and hence the values are either 0 or 1.

**FIG. 6:**
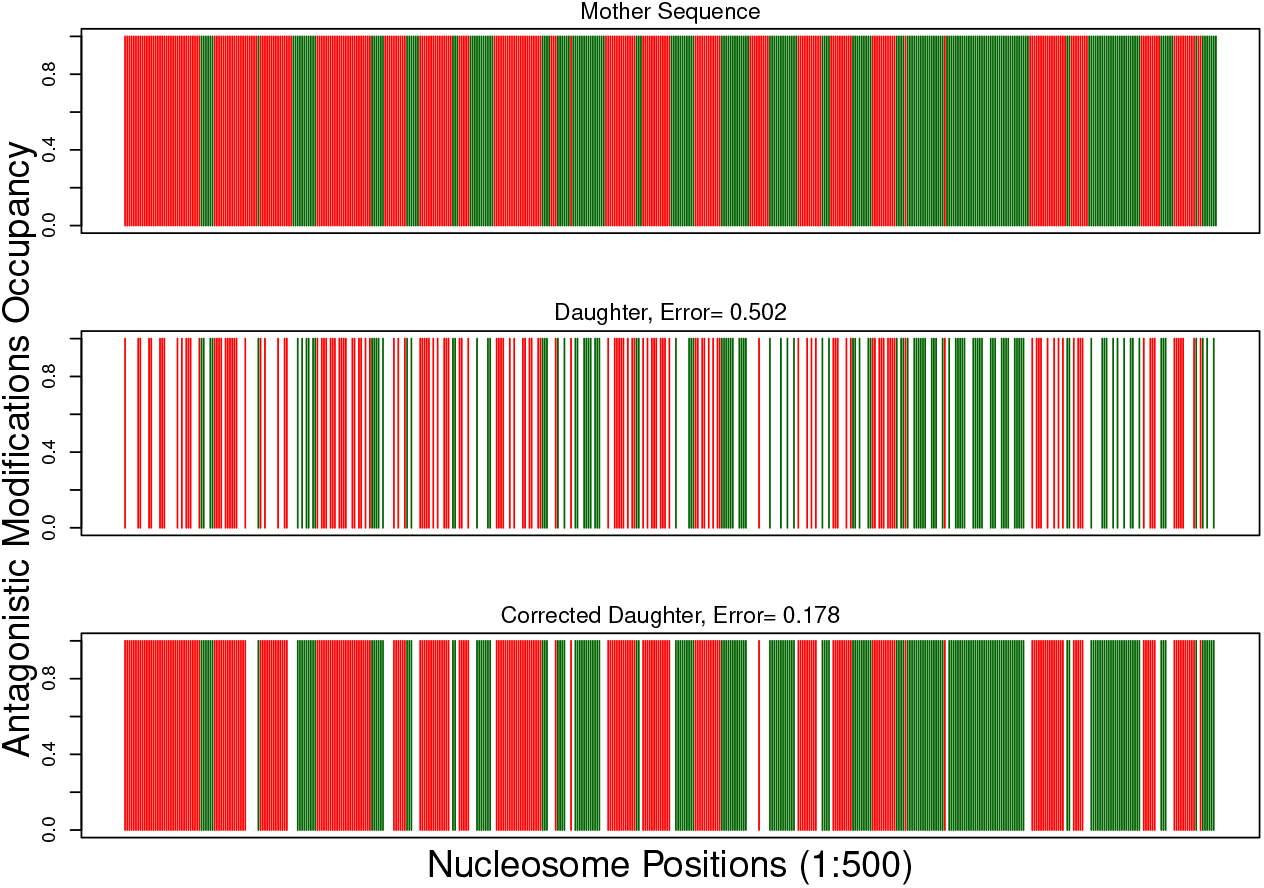
Simulating antagonistic modifications: Typical realizations of the modification patterns from the study of two spatially distinct/antagonistic modifications. The red and green regions represent the modifications 1 and 2, respectively, spread over 500 nucleosomes. Corrections were performed using the threshold-*k* algorithm with an optimum *k*_*t*_ = 6.

## DISCUSSION

In this work, we proposed that the problem of the daughter chromatin retrieving histone modification patterns, to achieve a mother-like chromatin state, can be mapped to a communication theory problem of receiving noisy signal and correcting it to retrieve the original signal. Using ideas from Information theory, we argued that if enzymes were ideal computing machines, the best they could do is to execute a MAP decoding algorithm to get back a mother-like sequence. We showed how well this algorithm would reconstruct the mother – the error can be as low as 5% in certain parameter regimes. However, the question was whether realistic enzymes can practically do such complex algorithms. We showed that in a biologically relevant parameter regime, MAP decoding algorithm is equivalent to a *k*-threshold filling algorithm. That is, the enzymes could simply insert modifications in *k*-sized or shorter unmodified stretches (0s). The fact that a detailed theory simplifies to a process that is potentially executable by enzymes makes this result attractive.

We modeled the mother chromatin as a first order Markov process. This is a reasonable model as there are minimal number of parameters (*α* and *β*) and the parameters are explicitly related to experimentally measurable properties of the modification patterns such as the mean contiguous length of modified and unmodified regions(see SI Sec. IV). By varying *k*_*t*_, as shown in FIGS. 4(B) and 5(A), we can determine the optimal configurations and obtain insights about how enzymes might work without a-priori knowledge of the statistical parameters. Note that even in the large *α* regime (region *a* in FIG. 4(A)), if enzymes settle for *k*-filling with a finite *k*_*t*_ (e.g., 5 or 6), it becomes a pragmatic modification correction solution as the resulting error is relatively low (see red and green curves in FIG. 4(C)).

It has to be noted that our model picks configurations (sequences) that maximise certain conditional probabilities, analogous to the minimum energy (or “equilibrium”) solution in physics. The model does not involve precise kinetic moves. There could be multiple different kinetic events leading to the same static configuration. Hence, the predictions of our model may be interpreted in terms of various known kinetic events in the field. For example, the determination of critical threshold *k*_*t*_ as well as the decision to fill the gaps would involve local read-write mechanisms as well as feedback loops. The decision to leave long gaps unfilled would also involve certain feedback mechanisms. For example, the effect of antagonistic modifications can be considered as a case of a negative feedback loops.

What might be the biological processes that keep a group of nucleosomes in a modified state or unmodified state is an interesting question. A region can be kept unfilled as a combined effect of modification and de-modification (de-methylation, de-acetylation etc.) events. When the rate of de-modification in a region is much higher than the modification rate, the region can remain unfilled. Physically, this could happen in several ways. Enzymes could cooperatively act on a small region. For example, interesting recent experimental studies [55] suggest that enzymes that act on histone modifications can phase separate to form a droplet around a group of nucleosomes, and may collectively modify/demodify all the nucleosomes together. It has also been shown that chromatin itself can form nano-domains having size of a few nucleosomes paving way for collectively maintaining a certain modification state in these groups of nucleosomes [56–58]. The potential enzymes/complexes that could do these activities could be JmjC domain proteins, UTX, NuRD, Fbxl10 and JARID1A [54, 59, 60]. Approaching this question from a different angle, currently, it is well-known that there exists certain correlations and anti-correlations between modifications [13]. There are biological mechanisms which are known to maintain these correlated states between modifications [33]. Hence, the mechanism that may cause the absence of a modification could also cause the presence of an opposite modification as we have discussed in the antagonistic modifications section. For example, recent experiments from [42] suggest that there is mutual antagonism between H3K27 and H3K36 methylation. This antagonism could also be one of the reasons for a long set of nucleosomes in which one modification is absent.

How enzymes know what is the optimal threshold for filling, is another interesting question. There are different families of enzymes, and each may have evolved differently. Some enzymes may be naturally adept at filling short regions, i.e., optimal *k* value (*k*^*^) could be hard-wired into such enzymes, via evolution. Another possibility is that other phenomena like local looping, phase separation etc. decide the threshold *k*^*^, by bringing unfilled nucleosomes together [61, 62]. These need to be understood in more detail in the future. As we showed, the information theory ideas suggest that the threshold-*k* filling algorithm emerges as a natural primary solution. However, the reconstruction may involve secondary mechanisms like boundary determination (e.g. CTCF [63]), synergistic modification between correlated modifications, and DNA methylation maintenance over the course of the cell-cycle [24, 33].

In this manuscript, we have been computing the deviation for every bit (BER). However, one can also quantify the deviation by other means; for example, using blocks of bits. That is, we take a small block of nucleosomes, find the average modification value and compare it with that of the equivalent block in the daughter cell, (see SI Sec VIII). In such a case, we find that the error decreases with increasing block size (FIG. S3). If the biological systems compute the errors in blocks of nucleosomal patterns, then this metric can be useful.

It is also interesting to discuss the threshold *k* − filling model in the context of existing statistical models. In a series of papers, Sneppen and collaborators [34, 54] suggest a bistable paradigm where the mean modification of a region decides its chromatin state. They showed that to achieve this bistable state, one requires long range interactions among nucleosomes. However, unlimited range of interaction can lead to spatially uniform pattern, which is not what is observed in experiments like [26]. It is in this context that our work is relevant; our finding of *k*-threshold filling bridges this gap and provides a natural length-scale (of length *k*_*t*_) for such long-range interactions. It is also important to note that in certain cell types (e.g. stem cells) when the nucleosomes are in a bivalent state, the bistable paradigm might be more relevant while in differentiated cells, where certain genes must always be kept in the repressed state, our paradigm of reconstructing a more precise pattern (as opposed to maintaining a mean modification state) could be more relevant.

Finally, it would also be of great interest to study how the polymer nature of the chromatin would work in tune with the results from information theory. After all, the epigenetic code might involve an interplay between the one-dimensional histone codes and polymer dynamics of the chromatin. The fact that we have two 1*s* at the boundaries suggests some potential role of looping or micro phase separation in far-away regions coming together. Our own earlier studies [58] hint to us that small patches of unmodified nucleosomes (newly inserted nucleosomes) may lead to small clusters, influencing the kinetics of the modification process itself. These are questions that await future studies.

## ACKNOWLEDGEMENTS

RP acknowledges funding from Department of Science and Technology India via Science and Engineering Research Board grant EMR/2016/ 005965, and Department of Biotechnology support via BT/HRD/NBA/39/12/2018-19.

## Conflict of Interests

The authors declare that they have no conflict of interests.

## Notes

### Competing Interest Statement

The authors have declared no competing interest.

