## Supplemental text for "High fidelity epigenetic inheritance: Information theoretic model predicts *k*-threshold filling of histone modifications post replication"

(Dated: May 23, 2021)

### I. MAXIMUM À-POSTERIORI PROBABILITY DECODING

As illustrated in FIG. 1(C) of the manuscript, the vector **M** represents the binary mother sequence, and each non-zero value of this sequence is independently flipped with probability half to obtain the daughter sequence **D**. This daughter sequence represents one of the two possible chromatins post replication. The flipping operation is equivalent to a logical AND between the independent sequences **M** and **Z**, where the **Z** is independently and identically distributed (IID) according to an unbiased binary random variable. The goal now is to restore a mother-like pattern $\hat{\mathbf{M}}$ from **D**. Recall the notation $m_i^j = m_i, m_{i+1}, \ldots, m_j$ introduced in the manuscript. Using this, let $m_1^N$ represent a realization of the mother sequence **M** of length $N$.

We use the MAP (Maximum À-posteriori Probability) rule to reconstruct a mother like sequence from the daughter. The MAP rule [1] suggests to choose the sequence $\hat{\mathbf{M}}$ which maximizes the à-posteriori probability $\mathbb{P}(\hat{\mathbf{M}}|\mathbf{D})$. Thus, the decoder chooses $\hat{\mathbf{M}}$ such that

$$\begin{aligned}\hat{\mathbf{M}} = \underset{\mathbf{M}}{\operatorname{argmax}}\ \mathbb{P}(\mathbf{M}|\mathbf{D}) &= \underset{\mathbf{M}}{\operatorname{argmax}} \left(\frac{\mathbb{P}(\mathbf{M})\mathbb{P}(\mathbf{D}|\mathbf{M})}{\mathbb{P}(\mathbf{D})}\right) \\ &= \underset{\mathbf{M}}{\operatorname{argmax}} \big(\mathbb{P}(\mathbf{M})\mathbb{P}(\mathbf{D}|\mathbf{M})\big).\end{aligned} \tag{S1}$$

The first step in Eq. S1 follows from the Bayes theorem. The second step uses the fact that $\mathbb{P}(\mathbf{D})$ is just a scaling factor while maximizing over **M**, and the argument which maximizes is unchanged by removing any common scaling.

The quantity $\mathbb{P}(\mathbf{M})$ in Eq. S1 is governed by Eq. (1) in the manuscript.

$$\mathbb{P}\big(\mathbf{M} = m_1^N\big) = \mathbb{P}(m_1)\prod_{i=2}^{N}\mathbb{P}(m_i|m_{i-1}). \tag{S2}$$

Taking $m_0 = \emptyset$ (empty set), we can write

$$\mathbb{P}\big(\mathbf{M} = m_1^N\big) = \prod_{i=1}^{N}\mathbb{P}(m_i|m_{i-1}). \tag{S3}$$

To prove Eq. (2) in the manuscript,

$$\begin{aligned}\mathbb{P}(\mathbf{D} = d_1^N | M = m_1^N) &= \mathbb{P}(d_1, d_2, \ldots, d_N\ |m_1^N) \\ &= \mathbb{P}(d_1 \mid m_1^N)\mathbb{P}(d_2 \mid m_1^N, d_1)\ldots\mathbb{P}(d_N \mid m_1^N, d_1^{N-1}) \\ &= \mathbb{P}(d_1|m_1)\mathbb{P}(d_2|m_2)\ldots\mathbb{P}(d_N|m_N) \\ &= \Pi_{i=1}^N\mathbb{P}(d_i|m_i).\end{aligned} \tag{S4}$$

The second step in Eq. S4 follows from Bayes rule. Since $d_i = m_i$ AND $z_i$, and the realization $z_i$ is independent of $(z_1^{i-1}, z_{i+1}^N, m_1^N)$ by our IID assumption on flipping, we have $\mathbb{P}(d_i|m_1^N, d_1^{i-1}) = \mathbb{P}(d_i|m_i)$. Notice that this corresponds to the memoryless nature of the flipping operation.

Thus Eq. (5) of the manuscript follows from equations (S3) and (S4).

---

*Electronic address:
†Electronic address:
‡Electronic address:

### II. MARKOV PROCESS, TRELLIS DIAGRAM AND BRANCH PROBABILITIES

By the definition of our modification flipping operation, we obtain

$$\mathbb{P}\big(d_i = 0 \mid m_i = 0\big) = 1 \tag{S5}$$

$$\mathbb{P}\big(d_i = 1 \mid m_i = 1\big) = \mathbb{P}\big(d_i = 0|m_i = 1\big) = \frac{1}{2}. \tag{S6}$$

The ideas behind trellis decoding is best illustrated with a generic example. Given a daughter sequence $d_1^N$, consider two possible mother sequences $a_1^N$ and $b_1^N$. The MAP rule will prefer sequence $a_1^N$ over $b_1^N$ if the joint probability law obeys $\mathbb{P}(a_1^N, d_1^N) > \mathbb{P}(b_1^N, d_1^N)$. The trellis diagram, depicted in FIG. 2 of the manuscript, is an effective way to compute and compare these joint probabilities. We can limit our considerations to sequences $a_1^N, b_1^N$ and $d_1^N$ which start and end with the value 1, since $d_1^N = (1, 0_{N-2}, 1)$ is given as the observed sequence. In the expressions below, joint probability terms of the form $\mathbb{P}(x, y|w)$ will have the first variable $x$ representing the mother and the second variable $y$ representing the daughter, at a particular nucleosome index.

$$\begin{aligned}\mathbb{P}(a_1^N, d_1^N) &= \mathbb{P}(1,1)\left(\prod_{i=2}^{N-1}\mathbb{P}(a_i, d_i = 0|a_{i-1})\right)\mathbb{P}(a_N, d_N = 1|a_{N-1})\\ &= \mathbb{P}(1,1)\left(\prod_{i=2}^{N-1}\mathbb{P}(a_i|a_{i-1})\mathbb{P}(d_i = 0|a_i)\right)\mathbb{P}(a_N|a_{N-1})\mathbb{P}(d_N = 1|a_N).\end{aligned} \tag{S7}$$

Similarly,

$$\mathbb{P}(b_1^N, d_1^N) = \mathbb{P}(1,1)\left(\prod_{i=2}^{N-1}\mathbb{P}(b_i|b_{i-1})\mathbb{P}(d_i = 0|b_i)\right)\mathbb{P}(b_N|b_{N-1})\mathbb{P}(d_N = 1|b_N). \tag{S8}$$

In the both the above equations we used the fact that $a_1 = b_1 = d_1 = d_N = 1$. Notice that, while comparing (S7) and (S8), we can ignore the initial common scaling factor $\mathbb{P}(1,1)$. The trellis diagram enables finding the remaining product by assigning branch metrics to the $N-1$ possible transitions of each path. In particular, the $i^{th}$ transition corresponding to the sequence $a_1^N$ will have an associated branch metric $\mathbb{P}(m_{i+1}|m_i)\mathbb{P}(d_{i+1}|m_{i+1})$ in the trellis diagram. From our Markov model, the probabilities $\mathbb{P}(m_i, d_i|m_{i-1})$ take the form

$$\mathbb{P}(0,0|0) = \beta \tag{S9}$$

$$\mathbb{P}(0,0|1) = 1-\alpha \tag{S10}$$

$$\mathbb{P}(1,0|0) = \mathbb{P}(1,1|0) = \frac{1-\beta}{2} \tag{S11}$$

$$\mathbb{P}(1,0|1) = \mathbb{P}(1,1|1) = \frac{\alpha}{2} \tag{S12}$$

$$\mathbb{P}(0,1|0) = \mathbb{P}(0,1|1) = 0. \tag{S13}$$

Let us now illustrate the trellis computations for a specific example. Take $N = 4$, and $a_1^4 = (1,1,1,1)$ and $b_1^4 = (1,0,0,1)$. The observed daughter sequence is given as $(d_1, d_2, d_3, d_4) = (1,0,0,1)$.

$$\mathbb{P}(a_1^4, d_1^4) = \mathbb{P}(1,1)\mathbb{P}(1,0|1)\mathbb{P}(1,0|1)\mathbb{P}(1,1|1) \tag{S14}$$

$$= \mathbb{P}(1,1)\frac{\alpha}{2}\frac{\alpha}{2}\frac{\alpha}{2} \tag{S15}$$

$$= \mathbb{P}(1,1)\,\frac{\alpha^3}{8}. \tag{S16}$$

$$\mathbb{P}(b_1^4, d_1^4) = \mathbb{P}(1,1)\mathbb{P}(0,0|1)\mathbb{P}(0,0|0)\mathbb{P}(1,1|0) \tag{S17}$$

$$= \mathbb{P}(1,1)\left((1-\alpha)\,\beta\,\frac{1-\beta}{2}\right) \tag{S18}$$

$$= \mathbb{P}(1,1)\,\frac{(1-\alpha)\beta(1-\beta)}{2}. \tag{S19}$$

Trellis decoding will compute $\frac{\alpha^3}{8}$ as the metric for the path $a_1^4$, whereas $\frac{1}{2}(1-\alpha)\beta(1-\beta)$ will be the metric of the path $b_1^4$. As another example, from FIG. 2 of the manuscript, the metric for the path $(1,0,0,0,0,0,1)$ can be easily computed by traversing through the corresponding path and reading out the product of branch metrics encountered, leading to a final metric $(1-\alpha)\beta^5\frac{1}{2}(1-\beta)$.

### III. SMAP ALGORITHM METHODOLOGY AND RESULTS

#### SMAP Decoding Proposition Proof

**Proposition 1** *Let $i, j$ be two positions where the daughter sequence has ones, with $j > i$. Then, SMAP decoding will choose a sequence having $\hat{m}_i^j$ according to*

$$\hat{m}_i, \ldots, \hat{m}_j = \textit{argmax}\, \mathbb{P}(m_i, \ldots, m_j | d_i = 1, d_{i+1}^{j-1}, d_j = 1).$$

Notice that $\mathbb{P}(m_i, d_i|m_{i-1}) = \mathbb{P}(m_i|m_{i-1})\mathbb{P}(d_i|m_i, m_{i-1}) = \mathbb{P}(m_i|m_{i-1})\mathbb{P}(d_i|m_i)$, by Bayes rule and Eq.(2) in the manuscript. Using this in Eq.(5) of the manuscript,

$$\mathbb{P}(\mathbf{M}|\mathbf{D}) = \frac{1}{\mathbb{P}(\mathbf{D})} \prod_{l=1}^{i-1} \mathbb{P}(m_l, d_l|m_{l-1})\mathbb{P}(m_i|m_{i-1})\mathbb{P}(d_i|m_i) \prod_{l=i+1}^{j} \mathbb{P}(m_l, d_l|m_{l-1}) \prod_{l=j+1}^{N} \mathbb{P}(m_l, d_l|m_{l-1}). \tag{S20}$$

Given the actual value of $m_j$, determining the bits $\hat{m}_{j+1}^N$ can be easily seen from above as

$$\hat{m}_{j+1}^N = \text{argmax} \prod_{l=j+1}^{N} \mathbb{P}(m_l, d_l|m_{l-1}). \tag{S21}$$

Similarly, once $m_i$ is fixed, then the bits $\hat{m}_1^{i-1}$ are determined by

$$\hat{m}_1^{i-1} = \text{argmax} \prod_{l=1}^{i-1} \mathbb{P}(m_l, d_l|m_{l-1})\mathbb{P}(m_i|m_{i-1}). \tag{S22}$$

Observe that both (S21) and (S22) are independent of $m_{i+1}^{j-1}$. Given $d_i = d_j = 1$, only sequences with $m_i = m_j = 1$ are to be considered for our maximization, since $\mathbb{P}(m_i = 1|d_i = 1) = 1$. Thus, the subsequences $\hat{m}_1^{i-1}$ and $\hat{m}_{j+1}^N$ are determined independently of $\hat{m}_{i+1}^{j-1}$, as shown above. Therefore we get

$$\hat{m}_{i+1}^{j-1} = \underset{m_i^j : m_i = m_j = 1}{\text{argmax}} \prod_{l=i+1}^{j} \mathbb{P}(m_l, d_l|m_{l-1}) \tag{S23}$$

as the output subsequence of SMAP decoding. The proof of the proposition is completed by noting that $\mathbb{P}(m_i^j|d_i^j) = \mathbb{P}(m_i^j, d_i^j)/\mathbb{P}(d_i^j)$ and $\mathbb{P}(m_i^j, d_i^j) = \prod_{l=i}^{j} \mathbb{P}(m_l, d_l|m_{l-1})$.

#### Computation of Statistical Properties

In this section, we explain how the parameters $\alpha$ and $\beta$ for a given binary sequence, modeled by the Markov process described in FIG. 1(b) of the manuscript, can be estimated. Consider a realization $\{m_1, m_2, \ldots, m_N\}$ of the binary sequence $\mathbf{M}$. We define the following variables as the number of transitions:

$$\begin{aligned} &n_{11} - \text{Number of transitions from 1 to 1} \\ &n_{10} - \text{Number of transitions from 1 to 0} \\ &n_{01} - \text{Number of transitions from 0 to 1} \\ &n_{00} - \text{Number of transitions from 0 to 0.} \end{aligned} \tag{S24}$$

While traversing through the sequence, we increment the transition variables as follows:

$$\begin{aligned} n_{11} &= n_{11} + 1;\ \text{if } (m_{i-1} = 1\ AND\ m_i = 1); \\ n_{10} &= n_{10} + 1;\ \text{if } (m_{i-1} = 1\ AND\ m_i = 0); \\ n_{01} &= n_{01} + 1;\ \text{if } (m_{i-1} = 0\ AND\ m_i = 1); \\ n_{00} &= n_{00} + 1;\ \text{if } (m_{i-1} = 0\ AND\ m_i = 0). \end{aligned} \tag{S25}$$

We then compute $\alpha$ and $\beta$ as:

$$\alpha = \frac{n_{11}}{(n_{11} + n_{10})} \tag{S26}$$

$$\beta = \frac{n_{00}}{(n_{00} + n_{01})}. \tag{S27}$$

The biological processes that keep nucleosomes in a modified state or unmodified state (leading to the parameters $\alpha$ and $\beta$ ) include co-operative modification/de-modification between correlated modifications, the effect of the presence of antagonistic modifications in certain nucleosomes, amongst others. When the rate of de-modfiication is high as compared with the rate of modification, longer runs of 0s can be observed. Some potential enzymes that perform de-modification (de-methylation, de-acetylation , de-ubiquitination) could be JmjC domain proteins and UTX , NuRD, Fbxl10 and JARID1A [2, 3]

#### Simulation Results

We simulated the mother sequences in a computer using different values of $\alpha$ and $\beta$ for the Markov process. Given a mother sequence, daughter sequences were generated by randomly flipping the non-zero values using independent realizations of an unbiased coin. 200 such daughter realizations were generated for each mother sequence considered. Each daughter sequence was corrected using the SMAP decoding rule, and compared to its mother to obtain the mean error ($\bar{\Delta}$). The error was averaged over all the 200 daughter sequences, and the whole experiment was repeated 300 times to obtain the averaged error (averaged over a total of 60000 cases) . The length of the sequence was taken as 100 nucleosomes in the simulations. The variation of the mean error across $\alpha$ and $\beta$ are plotted in FIG. S1. One could correlate FIG. 3 of the manuscript which has the heatmap of the mean error after correction and compare with these plots to understand the regions of low and high error. Furthermore, the behaviour shown in the plots correlate with the phase diagram FIG.4(A) of the manuscript.

### IV. ANALYSIS OF STATISTICAL PROPERTIES OF MODIFICATION DATA

#### Average number of 0s and 1s in the mother sequence

The steady state probabilities ($\Pi_1$ and $\Pi_0$) of states '1' and '0' in a binary sequence modelled by a Markov chain described in FIG. 1(b) of the manuscript are given by the following equations [4]:

$$\Pi_1 = \frac{(1-\beta)}{(2-\alpha-\beta)} \tag{S28}$$

$$\Pi_0 = \frac{(1-\alpha)}{(2-\alpha-\beta)}. \tag{S29}$$

The average number of 1s and 0s in the mother sequence is given by the steady state probabilities of states '1' and '0' in Eq. S28 and Eq. S31, multiplied by the length of the sequence.

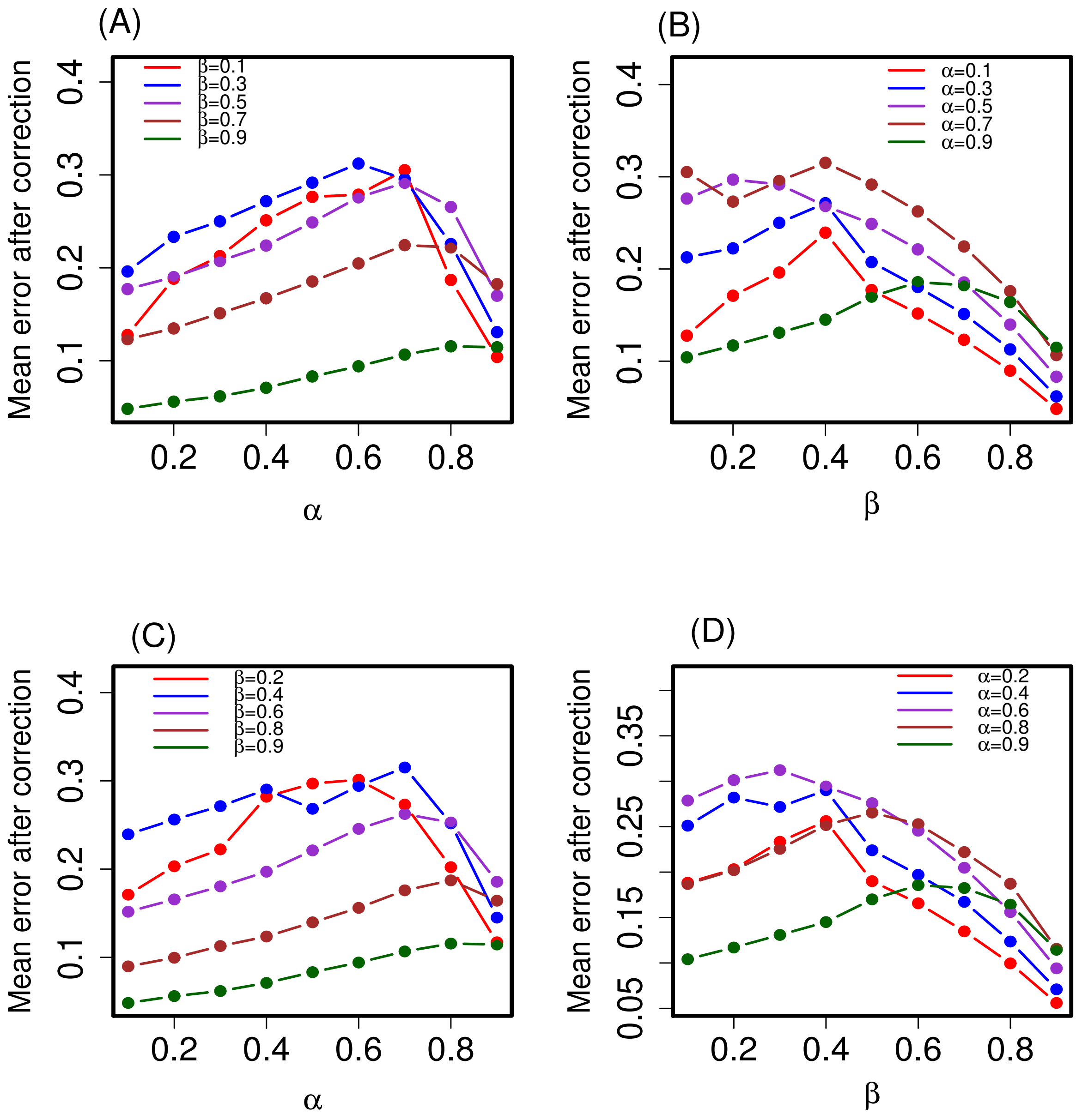


FIG. S1: The variation of the mean deviation after correction ($\bar{\Delta}$) using the SMAP algorithm is plotted as a function of different $\alpha$ and $\beta$ in the above figures. The error is averaged over a simulation of 300 mother sequences each of which has 200 daughter sequences (a total of 60000 cases). In (A) and (C), we plot the mean error after correction as a function of $\alpha$ for mostly (A) odd and (C) even values of $\beta$. In (B) and (D), we plot the mean error after correction as a function of $\beta$ for mostly (B) odd and (D) even values of $\alpha$. The error bars representing SEM are are smaller than the size of the points in the plot. It can be observed that in some regions of combinations of very high or very low $\alpha$ and $\beta$ , the mean error is quite low.

#### Average number of intermediate 0s and 1s

Let us compute the average number of intermediate 0s between two 1s in a long sequence of the mother sequence. Based on our Markov model, the run of zeros can be considered as a geometric random variable that lists the number of 0s before the next 1 is encountered. The probability of getting a 1 after a 0 is $1 - \beta$. Thus, given that there is at least one 0, the expected number of 0s between two 1s is the mean of this geometric random variable, given by

$$\text{Expected number of intermediate 0s} = \frac{1}{1-\beta}. \tag{S30}$$

In other words, this is the mean contiguous length of intermediate 0s.
Similarly, the average length of a run of 1s between two bordering 0s can be computed as

$$\text{Expected number of intermediate 1s} = \frac{1}{1-\alpha}. \tag{S31}$$

Both these quantities can be obtained from experimental data.
From Eq. S28, Eq. S31 and the values of the average number of intermediate 0s and 1s , we understand that when $\alpha$ and $\beta$ are very high, the average run of 0s (or 1s) will be high. This would result in high chances of finding a long stretch of 0s, or a long island of 1s, for large values of $\alpha$ and $\beta$.

### V. DISCRETIZATION ALGORITHM

The population-averaged parental H3K27me3 data obtained from the experiments in [5] were converted to binary realizations of a single-cell indicating the presence or absence of the H3K27me3 modification in a nucleosome. The algorithm is explained in the flowchart of FIG. S2. It has to be noted that the H3 experimental data was also used in the algorithm in order to detect the presence of the nucleosomes as against finding the presence of the H3K27me3 modification in the nucleosome. The files chosen for FIG. 5 in the manuscript are $GSM2988386\_H3K27me3\_ChIPseq\_Parental\_RPM\_rep1.bedgraph$ (H3K27me3 modification data) and $GSM2988395\_H3\_ChIPseq\_Parental\_RPM\_rep1.bedgraph$(control H3). Notice that we employed a naive discretization scheme, the primary purpose was to validate the regime of relevant $(\alpha, \beta)$. More sophisticated discretization schemes will not significantly change the regime.

### VI. EXPERIMENTAL DATA - METHODS AND RESULTS

#### Statistical Analysis of Experimental Data

To obtain the binary realizations of the H3K27me3 data from [5], we obtained the nucleosomes in the region of bp:151,495,060-165,790,665 and applied the discretization algorithm explained in FIG. S2. We obtained 100 binary realizations of the mother. The average values of $\alpha$ and $\beta$ for mother realizations were computed using (S26)–(S27) to yield 0.81 and 0.815 respectively. Each of the mother sequences were ANDed with a binary random binomial sequence (with bias 0.5) to produce 100 daughters. They were corrected with the threshold-k filling algorithm with different values of $k_t$. The results are plotted in FIG. 5 of the manuscript.

### VII. SIMULATION OF ANTAGONISTIC MODIFICATIONS

To simulate the spatially distinct antagonistic modifications illustrated in FIG. 6 of the manuscript, we adopted the following method. Since very high values of $\alpha$ and $\beta$ were known to produce long regions of 1s alternating with long regions of 0s, we used this approach to generate mother sequences with values 1 and 2 indicating the presence of modification-1 or modification-2. Daughter sequences were then obtained from these mother sequences by ANDing them with binary IID sequences to produce 0s indicating the absence of either modification.
While performing the corrections using a $k_t = 6$ with the threshold-$k$ filling algorithm, we filled those 0s which were bounded by 1s whose length was within threshold $k_t$ with 1s - simulating enzyme-1 for correcting modification-1. Similarly, we corrected those 0s which were bounded by 2s and whose length was within a threshold $k_t$ with 2s - simulating enzyme-2 for correcting modification-2. While a typical daughter sequence has an error $\sim$ 0.5, the corrected daughter had an error less than 0.2.

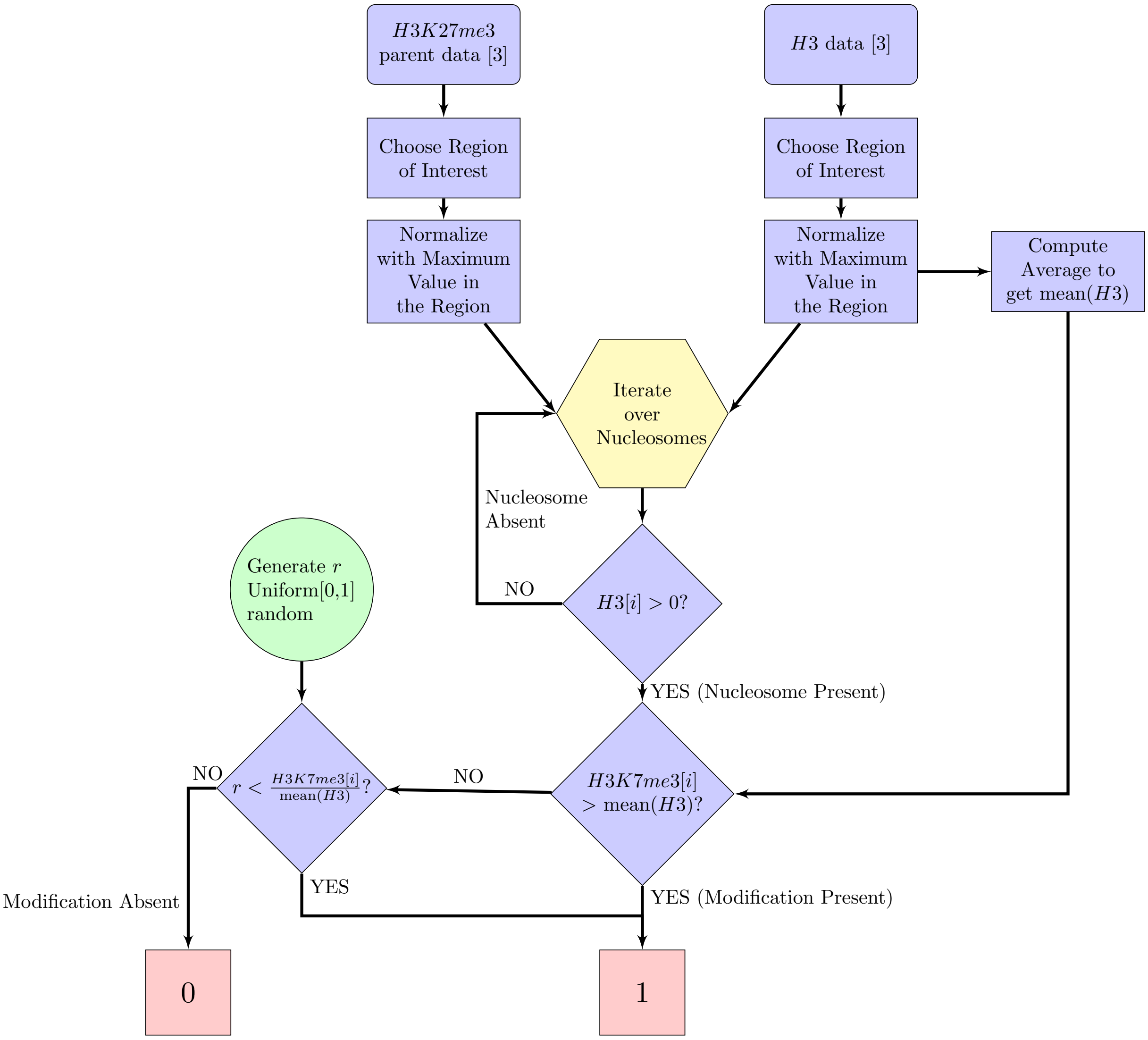


FIG. S2: The algorithm used to discretize the population-averaged parental H3K27me3 data from [5] into a binary realization - equivalent of a single cell representation indicating the presence (1) or absence (0) of the modification.

### VIII. BLOCK ERROR COMPUTATIONS

To account for the possibility that biological systems may be computing the similarity in patterns with the mother chromatin by comparing blocks of histone modifications, we computed the block error rate as follows: We averaged the bits over blocks of various sizes in a sliding window in both the mother chromatin and the corrected daughter chromatin. That is, in each array $\mathbf{M}$, starting from 1, a block of size $b$ nucleosomes were considered. The average of $b$ bits were computed as

$$\bar{m}_i = \frac{1}{b}\sum_{j=i}^{i+b} m_j$$

This process was repeated for the corrected daughter. The block averaged sqeuences were then used for computation of error using Eq. (3) in the manuscript. We computed the mean block error for the corrected sequences for different values for $k_t$ and plotted the results in FIG S3. The $\alpha$ and $\beta$ values of the generated mother sequences were 0.9 each and the length of the sequences is 100. One can observe that as the block size increases, the error decreases monotonically.

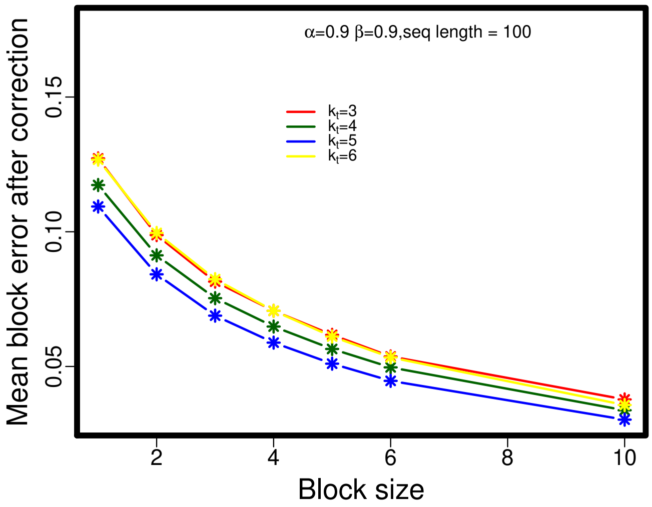


FIG. S3: The plot of the mean error computed in different blocks of nucleosomes between the original mother sequence and the daughter sequences for $\alpha$ and $\beta$ values of 0.9 each and $k_t = 3, 4, 5, 6$. With increasing block size, the mean error decreases monotonically.

[1] Cover TM, Thomas JA. 2012 *Elements of information theory.* John Wiley & Sons.
[2] Sneppen K, Ringrose L. 2019 Theoretical analysis of Polycomb-Trithorax systems predicts that poised chromatin is bistable and not bivalent. *Nature communications* **10**, 1–18.
[3] Swygert SG, Peterson CL. 2014 Chromatin dynamics: interplay between remodeling enzymes and histone modifications. *Biochimica et Biophysica Acta (BBA)-Gene Regulatory Mechanisms* **1839**, 728–736.
[4] Stewart WJ. 2009 *Probability, Markov chains, queues, and simulation: the mathematical basis of performance modeling.* Princeton university press.
[5] Reverón-Gómez N, González-Aguilera C, Stewart-Morgan KR, Petryk N, Flury V, Graziano S, Johansen JV, Jakobsen JS, Alabert C, Groth A. 2018 Accurate recycling of parental histones reproduces the histone modification landscape during DNA replication. *Molecular Cell* **72**, 239–249.
